# Restricted sequence variation in *Streptococcus pyogenes* penicillin binding proteins

**DOI:** 10.1101/2020.01.24.919308

**Authors:** Andrew Hayes, Jake A. Lacey, Jacqueline M. Morris, Mark R. Davies, Steven Y.C. Tong

## Abstract

A recent clinical report has linked *Streptococcus pyogenes* β-lactam antibiotic resistance to mutations in the Penicillin Binding Protein PBP2x. To determine whether this is an isolated case or reflects a broader prevalence of mutations that might confer reduced β-lactam susceptibility, we investigated the relative frequency of penicillin binding protein (PBP) sequence variation within a global database of 9,667 *S. pyogenes* isolates. We found that mutations in *S. pyogenes* PBPs (PBP2x, PBP1a, PBP1b and PBP2a) occur infrequently across this global database with less than 3 amino acid changes differing between >99% of the global population. Only 4 of the 9,667 strains contained mutations near transpeptidase active sites. The reported PBP2x T553K substitution was not identified. These findings are in contrast to those of 2,520 *S. pneumococcus* sequences where PBP mutations are relatively frequent and are often located in key β-lactam binding pockets. These data, combined with the general lack of penicillin resistance reported in *S. pyogenes* worldwide, suggests that extensive, unknown, constraints restrict *S. pyogenes* PBP sequence plasticity. These findings imply that while heavy antibiotic pressure may select for mutations in the PBPs, there is currently no evidence of such mutations becoming fixed in the *S. pyogenes* population nor that mutations are being sequentially acquired in the PBPs.

**Importance:** Penicillin is the first line therapeutic option for *Streptococcus pyogenes* infections. Despite the global high prevalence of *S. pyogenes* infections and widespread use of penicillin, reports of resistance to penicillin have been incredibly rare. Recently, penicillin resistance was detected in two clinical *S. pyogenes* isolates with accompanying mutations in the active site of the penicillin binding protein PBP2x, raising concerns that penicillin resistance may become more widespread. We screened a global database of *S. pyogenes* genome sequences to investigate the frequency of penicillin binding protein (PBP) mutations, identifying that PBP mutations are uncommon relative to *Streptococcus pneumoniae*. These findings support clinical observations that penicillin resistance is rare in *S. pyogenes*, and suggest that there are considerable constraints on *S. pyogenes* PBP sequence variation.

## INTRODUCTION

*Streptococcus pyogenes* (Group A *Streptococcus*, GAS) has previously been understood to be uniformly susceptible to β-lactam antibiotics (1). Two *S. pyogenes* isolates with elevated minimum inhibitory concentrations (MIC) to β-lactam antibiotics have recently been reported (2). Both isolates were molecularly typed as *emm* 43.4 and had a penicillin binding protein PBP2x missense mutation (T553K) at the transpeptidase active site which was associated with an 8-fold and 3-fold increased MIC to ampicillin and cefotaxime respectively compared to closely related isolates without the PBP2x mutation. In contrast to *S. pyogenes*, reduced susceptibility to β-lactams has been widely reported in *S. pneumoniae* and is strongly associated with sequence variation in PBPs (3, 4). Using GAS genome sequences from global sources, we sought to determine the prevalence of substitutions across the transpeptidase domains of the GAS PBPs (PBP2x, PBP1a, PBP2a and PBP1b) in comparison with *S. pneumoniae* (which shares PBP2x and PBP1a).

## METHODS

We obtained publicly available genome sequence data for 9,667 *S. pyogenes* isolates from the short-read archive (**Supplementary Table 1**). We assembled genomes using shovill v.1.0.9 (https://github.com/tseemann/shovill) with an underlying SKESA v.2.3.0 assembler (5). Using the β- lactam susceptible *S. pyogenes* serotype M3 strain ATCC BAA-595/MGAS315 as a reference, we determined the presence, amino acid sequence and alignment (6) of each of PBP2x, PBP1a, PBP1b and PBP2a in each genome with the screen_assembly script (7) and BlastP parameters of 100% coverage and 90% identity.

**Table 1:** Percentage of transpeptidase sequences with variation in the SXXK, SXN or K(T/S)G motifs of the transpeptidase active sites in PBP1a and PBP2x for *S. pneumoniae* and *S. pyogenes*.

|  | *S. pneumoniae* | *S. pyogenes* |
| --- | --- | --- |
| Motif | (n=2,520) | (n=9,667) |
|  | Variants (%) | Variants (%) |
| **<u>PBP1a</u>** |  |  |
| STMK* | 445 (17.7) | 0 (0) |
| SRN | 0 (0) | 0 (0) |
| KTG(T) | 0 (0) | 0 (0) |
| **<u>PBP2x</u>** |  |  |
| STMK* | 639 (25.3) | 4 (0.04) |
| SSN | 0 (0) | 0 (0) |
| KSG(T)** | 0 (3) (0 (0.1)) | 0 (0) |

\* the transpeptidase domain sequences as defined in Li et al 2016 were truncated between the S and T of the S//TMK
\*\* This table does not include the two T553K sequences reported to be associated with β-lactam resistance in *S. pyogenes* in Vannice et al.

To compare the conservation of the transpeptidase active site motifs across streptococcal species, full length PBP2x protein sequences of *S. pyogenes* serotype M3 strain ATCC BAA-595/MGAS315 (NC_004070.1), *S. pneumoniae* strain ATCC BAA-255/R6 (NC_003098.1), *S. agalactiae* strain 2603V/R (NC_004116.1), and the *S. dysgalactiae* subspecies *equisimilis* (SDSE) strain RE378 (NC_018712.1) reference genomes were aligned using Clustal Omega (8, 9). The percentage sequence similarity was compared using Blosum62 with threshold 1 in Geneious Prime (10).

To investigate the inferred crystal structure location of *S. pyogenes* PBP2x mutations relative to the *S. pneumoniae* orthologue, *S. pyogenes* PBP2x sequence variations were plotted onto the *S. pneumoniae* PBP2x crystal structure bound to oxacillin (PDB: 5OIZ) (11). Sequence conservation as determined by the frequency (for *S. pyogenes*) and percentage (for *S. pneumoniae*) of variant amino acids compared to the consensus was rendered onto the PBP2x crystal structure using UCSF Chimera (12).

We defined the PBP2x and PBP1a transpeptidase regions as that used in an assessment of 2,520 invasive *S. pneumoniae* isolates by Li et al (3) and determined and plotted the number of pairwise amino acid differences within these regions using Distances Matrix in Geneious Prime (10) and ggplot2 in R version 3.6.1 (13). Similarly, we also assessed the conservation of PBP1b and PBP2a proteins for the 9,667 *S. pyogenes* genomes, and the transpeptidase region of PBP2b for *S. pneumoniae*.

## RESULTS

We collated 9,667 *S. pyogenes* genome sequences, representing 115 different *emm* types and 321 multi-locus sequence types (**Supplementary Table 1, Supplementary Files 1-4**). These genome sequences were mostly from datasets from the United Kingdom and United States that focused on invasive disease (14-24).

Mutations in the penicillin binding proteins (PBPs) have been associated with reduced β-lactam susceptibility for *S. pneumoniae* (3), *S. agalactiae* (25), *S. dysgalactiae* (26), and now *S. pyogenes* (2). A comparison of PBP2x between β-lactam susceptible reference genomes of *S. pyogenes, S. pneumoniae, S. agalactiae*, and SDSE demonstrated a high level of inter-species conservation (>72% similarity, **Supplementary Figure 1 and Supplementary Table 2**). In *S. pneumoniae*, substitutions at the PBP2x transpeptidase active site (SXXK, SXN, and KSTG) result in reduced β-lactam susceptibility. These three motifs were conserved across the four species (**Supplementary Figure 1**).

**Figure 1:**
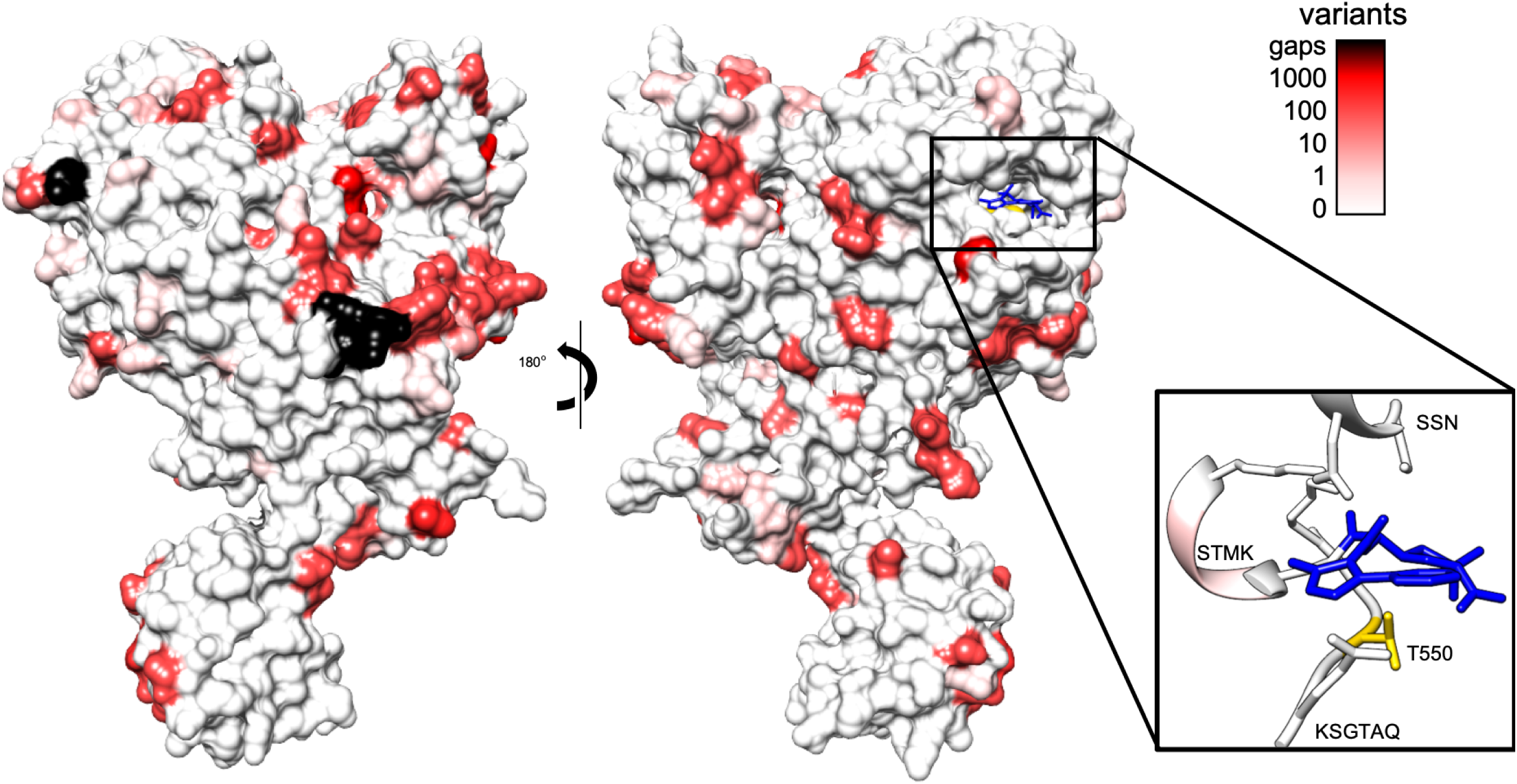
Global amino acid variation of *Streptococcus pyogenes* PBP2x mapped against the crystal structure of *Streptococcus pneumoniae* PBP2x. Crystal structure of PBP2X from *S. pneumoniae* bound to oxacillin (Blue) with residue conservation of *S. pyogenes* PBP2x mapped to surface. 9,667 *S. pyogenes* sequences were mapped to the 5OIZ structure sequence and frequency of conservation amongst *S. pyogenes* strains determined. Conservation was then mapped to surface residues and colour gradient applied. Black residues represent regions absent in the alignment due to absence of sequence relative to the *S. pneumonia* crystal structure. Thresholds were chosen to represent differing orders of magnitude for conservation with thresholds set at orders of magnitude (0, 1, 10, 100, 1000 sequences varying at the residue**). Inset**: ribbon diagram of binding pocket motifs SSN, STMK and KSG with position of mutated residue (T553K) highlighted (yellow). Mutations were observed in the STMK motif in 4 of the 9,667 sequences.

Given the similarity between PBP2x of *S. pneumoniae* and *S. pyogenes* (73.4% similarity, **Supplementary Table 1**), we mapped the conservation of residues from the alignment of 9,667 *S. pyogenes* PBP2x onto the crystal structure of *S. pneumoniae* PBP2x (**Figure 1**). The transpeptidase active site motifs are SXXK at positions 340-343 in *S. pyogenes* (positions 337-340 in *S. pneumoniae*), SXN at positions 399-401 in *S. pyogenes* (positions 395-397 in *S. pneumoniae*), and KSGT at positions 550-553 in *S. pyogenes* (positions 547-550 in *S. pneumoniae*). There were 105 unique amino acid sequence variants of the *S. pyogenes* PBP2x sequence with no major frameshifts or premature stop codons (**Supplementary File 1**). We found no instances of the T553K substitution in the PBP2x KSGT motif as reported in the recent *S. pyogenes* β-lactam resistant isolates (2). Only four *S. pyogenes* isolate sequences (0.04%) had substitutions within the transpeptidase active site motifs of PBP2x (**Figure 1** and **Table 1**) corresponding to STMK to SAMK and STMK to STIK. These changes may not have a phenotypic effect on penicillin susceptibility as STIK has been recently reported in a penicillin-susceptible isolate (GASAR0057) (17). Furthermore, no amino acid substitutions were found in the active site motifs of *S. pyogenes* PBP1a. In comparison, using population data from Li et al (3), *S. pneumoniae* had active site motif variants in 639/2,520 (25.3%) isolates for PBP2x and 445/2,520 (17.7%) for PBP1a (**Table 1**). A large proportion of *S. pneumoniae* substitutions mapped to areas near to the active site (**Supplementary Figure 2**).

**Figure 2:**
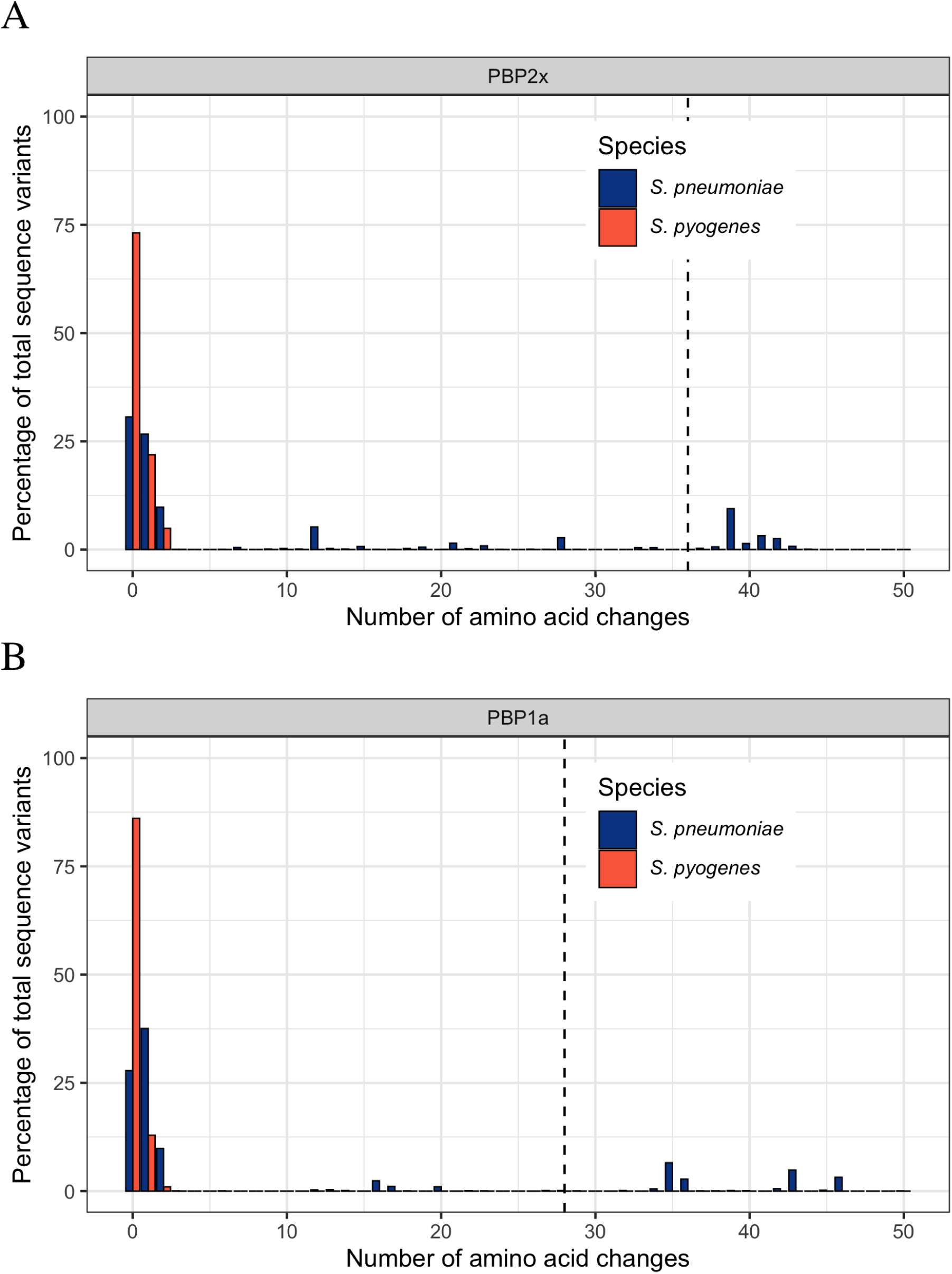
Amino acid differences of the transpeptidase domains of PBP2x and PBP1a. The percentage of isolates with changes in the transpeptidase domains of A) PBP2x and B) PBP1a, relative to penicillin susceptible references in *Streptococcus pneumoniae* (blue, n= 2,520) and *S. pyogenes* (red, n= 9,667). Sequences that are >10% divergent (indicated by dotted vertical lines) have been associated with increased penicillin minimum inhibitory concentrations in *S. pneumoniae*.

For *S. pneumoniae*, the number of substitutions across the whole transpeptidase domain of PBPs has been associated with penicillin resistance. Li et al. (3) found that penicillin MICs increased as the total number of divergent (defined as >10% amino acids different) transpeptidase domains of PBP2x, PBP1a and PBP2b increased from 0 to 3. For *S. pyogenes* we used the most common amino acid sequences of PBP2x and PBP1a as our reference and for *S. pneumoniae* a previously defined wildtype as the reference (3). There were considerably fewer PBP2x and PBP1a transpeptidase domains with multiple substitutions for *S. pyogenes* compared to *S. pneumoniae* (**Figure 2**). No *S. pyogenes* strains had sufficient mutations to reach the 10% threshold. For *S. pneumoniae*, 18.3% (462 of 2,520 strains) and 19.2% (485 of 2,520 strains) contained divergent PBP2x and PBP1a transpeptidase domains respectively (**Figure 2**). This pattern of greater conservation of *S. pyogenes* PBPs was also observed for PBP1b and PBP2a in *S. pyogenes* compared to PBP2b in *S. pneumoniae* (**Supplementary Figure 3**).

## DISCUSSION

We found no evidence that mutations are present in the β-lactam binding site KSGTAQ motif of PBP2x among 9,667 *S. pyogenes* genome sequences. Only four isolates contained mutations in the transpeptidase active sites of PBP2x and PBP1a. Although the report of two *S. pyogenes* isolates with reduced β-lactam susceptibility associated with *pbp2x* mutations is concerning (2), our findings provide reassurance that these are extremely limited, and perhaps unique, occurrences at this stage. We found a high degree of conservation of GAS PBP2x and PBP1a at transpeptidase active sites and across the broader transpeptidase domains. In comparison, PBP2x and PBP1a for *S. pneumoniae* were far less conserved, suggesting that there are strong evolutionary constraints in these domains for *S. pyogenes* that is not the case for *S. pneumoniae*. Studies of penicillin-resistant *S. pyogenes* generated through mutagenesis (27) or serial passage in penicillin containing medium (28), demonstrated that mutants with raised penicillin MICs appeared to have alterations in PBPs with reduced penicillin affinity (27). Notably mutants grow more slowly, have aberrant colony morphology compared to wild type strains (27), and are avirulent with a decrease in M protein production (28). These laboratory experiments, together with the absence of naturally occurring isolates with greater than five amino acid substitutions in PBP2x or PBP1a, strongly suggest that changes to the PBPs are associated with a significant fitness cost.

## ACKNOWLEDGEMENTS

This work was supported by an Australian National Health and Medical Research Council (NHMRC) project grant #1098319. SYCT is a NHMRC Career Development Fellow (#1145033). MRD is the recipient of a University of Melbourne C. R. Roper Fellowship. The funders had no role in study design, data collection and interpretation, or the decision to submit the work for publication.

---

**Supplementary Table 1:** Strain list with *emm* and MLST types, amino acid sequence and alleles of PBP2x, PBP1a, PBP1b and PBP2a.

**Supplementary Table 2:** The similarity matrix between PBP2x for four *Streptococcus* species as determined by BLOSUM62 threshold ≥1.

**Supplementary Figure 1: PBP2x alignment for *S. pyogenes, S. pneumoniae, S. agalactiae*, and *S. dysgalactiae subspecies equisimilis***

Clustal Omega multiple sequence alignment of PBP2x between of *S. pyogenes* serotype M3 strain ATCC BAA-595/MGAS315 (genome reference NC_004070.1, protein reference WP_011106648.1), *S. pneumoniae* strain ATCC BAA-255/R6 (genome reference NC_003098.1, protein reference NP_357898.1), *S. agalactiae* strain 2603V/R (genome reference NC_004116.1, protein reference NP_687322.1), and the *S. dysgalactiae subspecies equisimilis* (labelled *S. equisimilis*) strain RE378 (genome reference NC_018712.1, protein reference WP_015017311.1). The three transpeptidase active site motifs are highlighted in bold text and underlined. Fully conserved amino acids are denoted by an asterix, a strongly conserved protein is denoted by a colon and proteins that are weakly conserved are denoted by a full-stop.

**Supplementary Figure 2: *Streptococcus pneumoniae* transpeptidase domain sequences mapped to the crystal structure of PBP2x**

Crystal structure of PBP2X from *S. pneumoniae* bound to oxacillin (Blue) with residue conservation of the 118 transpeptidase variants identified by Li et al (2016) mapped to surface. The 118 non-redundant transpeptidase sequences identified by Li et al (2016) were aligned using MUSCLE aligner and the conservation of sites mapped to the surface of PDB 5OIZ and colour gradient applied. Black residues represent regions absent in the alignment due to not being part of the transpeptidase domain. Thresholds were chosen to represent the range of sequence variation in the unique sequences but unlike the *S. pyogenes* dataset does not represent frequency of the variants within the population. **Inset**: ribbon diagram of binding pocket motifs SSN, STMK and KSG with position of mutated residue (T550K) highlighted.

**Supplementary Figure 3:** Amino acid differences of the complete PBP1b and PBP2a for *S. pyogenes* and the transpeptidase domain of PBP2b for *S. pneumoniae*. The length of the complete PBP1b was 766 amino acids and PBP2a was 756 amino acids for *S. pyogenes*. The PBP2b transpeptidase domain was 280 amino acids in *S. pneumoniae*. The percentage of isolates with changes relative to penicillin susceptible references in the full protein of *S. pyogenes* (red; n= 9,667) and in the transpeptidase domains of *S. pneumoniae* (blue; n=2,520). For A) PBP1b (*S. pyogenes*), B) PBP2a (*S. pyogenes*) and C) PBP2b (*S. pneumoniae*). Sequences that are >10% divergent are indicated by dotted vertical lines.

